# Benchmarking Encoding Strategies for Non-Coding Mutations in Sequence-Based Prediction Tasks

**DOI:** 10.1101/2025.01.01.631025

**Authors:** Zhe Liu, Yihang Bao, Wenhao Li, Chengyi Yang, Weidi Wang, Wenxiang Cai, Weihao Li, Guan Ning Lin

**Author notes:** Authors to whom correspondence should be addressed. (Guan Ning Lin). These authors contributed equally to this work (Zhe Liu, Yihang Bao, Wenhao Li).

## Abstract

Non-coding single nucleotide polymorphisms (SNPs) are key modulators of gene regulation and have been implicated in diverse complex traits and diseases. With the growing demand for accurate functional interpretation of non-coding variants, the choice of encoding strategies becomes critical in downstream predictive modeling. Despite recent advances, a systematic evaluation of encoding approaches tailored for non-coding SNPs remains lacking. To address this gap, we present a comprehensive benchmark that evaluates six representative encoding strategies, including categorical, semantic, and functional embeddings, across three quantitative trait loci (QTL)-related prediction tasks. The study encompasses nine machine learning and deep learning models and incorporates experimental controls and repeated trials to ensure robustness and reproducibility. We assess each strategy along multiple dimensions, such as interpretability, representation abundance, and computational efficiency. Rather than ranking individual methods, our analysis emphasizes the interaction between encoding strategies, model types, and preprocessing protocols, and highlights their collective influence on predictive performance. This work establishes a standardized framework for evaluating non-coding SNP representations and offers actionable guidance for selecting and optimizing prediction pipelines in regulatory genomics.

## Introduction

Non-coding single nucleotide polymorphisms (SNPs) are critical genomic variants that influence gene regulation and phenotypic diversity without altering protein sequences[1-3]. These variants play key roles in regulating transcription, splicing, and epigenetic modifications, making them central to understanding disease mechanisms and genetic architecture[4, 5]. Despite their importance, decoding the functional impact of non-coding SNPs remains a challenging task due to their complex regulatory mechanisms and the subtle nature of their effects[6, 7].

Advances in genomic datasets, such as expression quantitative trait loci (eQTL)[8] and methylation quantitative trait loci (meQTL)[9], have created unprecedented opportunities to investigate the regulatory roles of non-coding SNPs. eQTL datasets elucidate how SNPs influence gene expression, while meQTL datasets provide insights into their effects on DNA methylation, an essential epigenetic modification. These datasets enable researchers to explore multiple layers of SNP-mediated regulation. However, the complexity of these tasks, combined with the diverse properties of available datasets (e.g., sample size, tissue specificity), underscores the need for systematic evaluation to identify effective encoding strategies and modeling approaches.

Computational tools have been developed to predict these impacts by encoding non-coding SNPs into machine-readable formats[10-13], yet the relative strengths and limitations of various encoding strategies and modeling approaches remain underexplored. A crucial step in addressing this challenge is exploring how to encode non-coding SNPs into features that computational models can process effectively. However, there is currently no objective evaluation or comprehensive guideline for selecting or designing encoding strategies tailored to sequence-based downstream prediction tasks. This lack of clarity leaves researchers uncertain about which encoding strategies are most effective for specific tasks and under what conditions they perform optimally. Bridging this gap requires a systematic benchmark to compare different encoding strategies and assess their applicability in modeling non-coding SNPs.

Various encoding strategies have been developed to transform non-coding SNPs into machine-readable features, broadly categorized into categorical, functional, and semantic strategies. Categorical strategies, such as One-Hot encoding, provide simple yet explicit representations of base-level information. Functional strategies, exemplified by models like Enformer[14], integrate genomic annotations to encode regulatory features. Semantic strategies, including pre-trained DNA language models like DNABert2[15], Genomic Pre-trained Network (GPN)[16], HyenaDNA[17], and Nucleotide Transformer (NT)[18], leverage deep learning to capture context-dependent relationships within DNA sequences. Each approach has distinct advantages and trade-offs, but their relative effectiveness in predicting downstream outcomes of non-coding SNPs, such as gene expression or DNA methylation changes, remains underexplored.

In this work, we present a comprehensive benchmark study to evaluate six distinct DNA sequence-based encoding strategies, ranging from traditional categorical encodings to advanced pre-trained semantic models and functional annotation-based embeddings. These strategies are assessed across key downstream tasks, including eQTL and meQTL prediction, using both machine learning and deep learning models. By systematically analyzing the impact of encoding strategies, model types, data sizes, and preprocessing strategies, we address critical questions about how to design prediction pipelines for non-coding SNPs. Our work emphasizes the need for clear guidelines in selecting encoding strategies and provides actionable insights into optimizing performance across different genomic contexts.

## Results

### Overview of benchmark framework

Non-coding single nucleotide polymorphisms (SNPs) play a pivotal role in gene regulation and disease mechanisms. However, the optimal strategies for encoding non-coding SNPs in computational downstream tasks remain unclear. To address this, we designed a benchmarking framework (Figure 1) that seeks to answer three fundamental questions:

**Figure 1.**
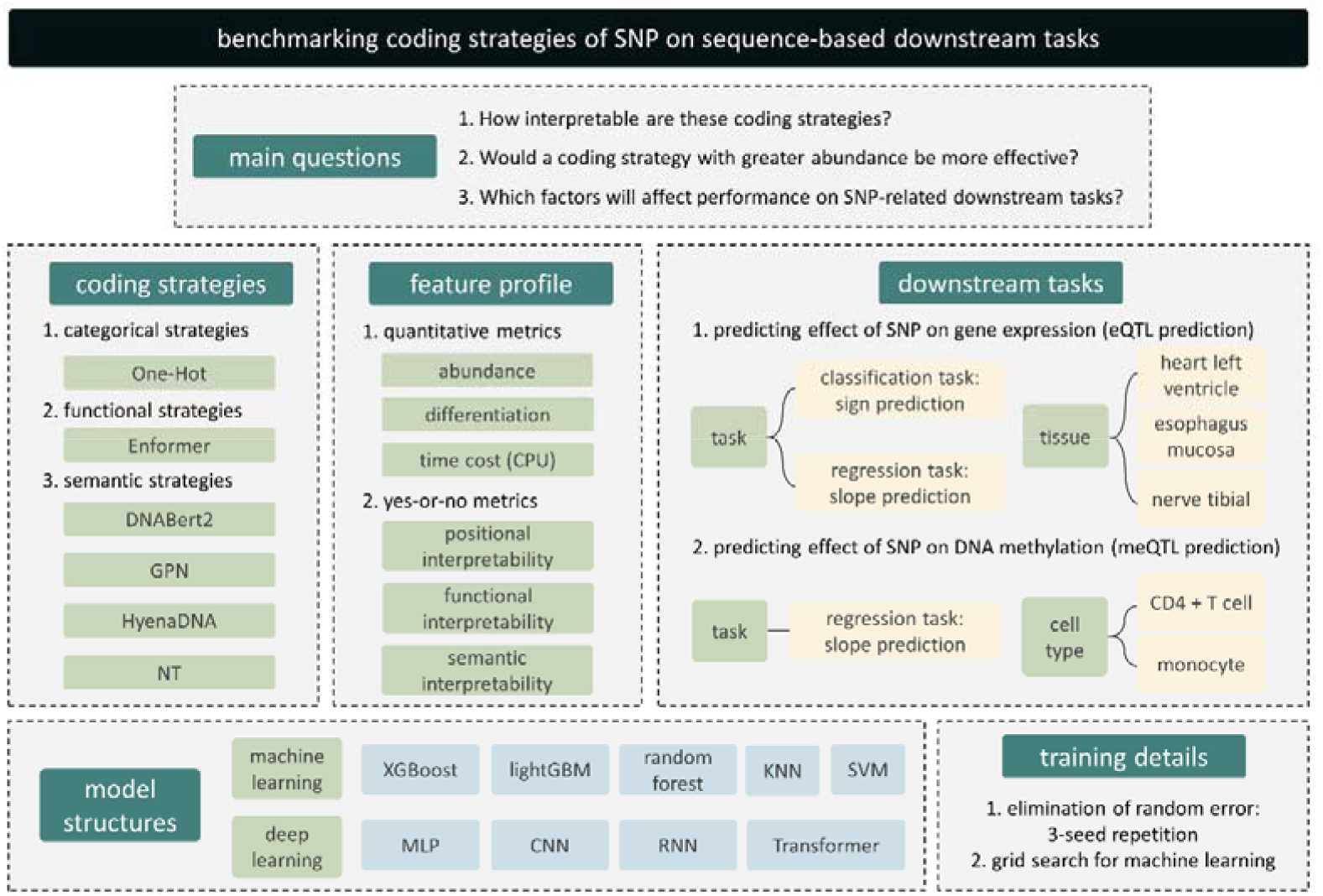
Framework for benchmarking coding strategies of non-coding SNPs. This framework evaluates six distinct coding strategies, categorized into categorical, functional, and semantic strategies. We designed six quantitative metrics to intuitively compare these strategies and assessed their performance across three Quantitative Trait Loci (QTL)-related downstream tasks. During the evaluation, a controlled variable approach was employed, with key variables such as coding strategies, machine learning/deep learning frameworks, tissue/cell types, data volume, and preprocessing strategies systematically observed.

1. How interpretable of these encoding strategies?
2. Are encoding strategies with higher encoding abundance (representation dimensions per base) more effective?
3. What are the key factors influencing performance in downstream tasks?

Our approach categorized six widely used encoding strategies into three conceptual groups: categorical, functional, and semantic strategies. These include the traditional One-Hot encoding method, the state-of-the-art sequence functional annotation representation model Enformer[14], and four pre-trained DNA language models: DNABert2[15], Genomic Pre-trained Network (GPN)[16], HyenaDNA[17], and Nucleotide Transformer (NT)[18]. These strategies were selected to represent a spectrum of encoding philosophies, ranging from simple base representation to complex contextual embeddings.

To systematically compare these strategies, we defined six key metrics, or feature profiles, aimed at quantifying their strengths and limitations. These include the ability to differentiate SNP effects, computational cost, compatibility with pooling mechanisms for sequence aggregation, and three complementary dimensions of interpretability: contextual (sequence-level), functional (annotation-level), and semantic (biological insight-level). Together, these metrics provide a multidimensional perspective on how encoding strategies translate genomic information into predictive tasks.

We applied this framework to three Quantitative Trait Loci (QTL)[19, 20]-related downstream tasks, investigating how non-coding SNPs influence gene expression (eQTL prediction, classification and regression tasks) and DNA methylation levels (meQTL prediction, regression task). These tasks spanned diverse tissue and cell types, incorporating both classification and regression objectives to ensure a holistic evaluation. Importantly, we considered not only the encoding strategies themselves but also the interactions between these strategies and model architectures. To this end, we tested a range of machine learning models, including eXtreme Gradient Boosting (XGBoost)[21], Light Gradient Boosting Machine (LightGBM)[22], Random Forest (RF)[23], k-Nearest Neighbors (KNN)[24], and Support Vector Machine (SVM)[25]. Additionally, we incorporated deep learning models such as Multi-Layer Perceptrons (MLP)[26], Convolutional Neural Networks (CNN)[26], Recurrent Neural Networks (RNN)[27], and Transformer-based architectures[28] to explore the capacity of these frameworks for representation learning.

To ensure robust conclusions, we controlled for potential confounding factors. All experiments were repeated using three different random seeds to reduce variability, while hyperparameters for machine learning models were optimized via grid search to achieve locally optimal performance. This rigorous experimental design not only enabled fair comparisons between encoding strategies but also allowed us to uncover key determinants of model performance, such as sample size, preprocessing methods, and model complexity.

By integrating a diverse set of methods, tasks, and experimental conditions, this benchmarking framework offers a comprehensive evaluation of encoding strategies for non-coding SNPs. It provides actionable insights into how these strategies can be tailored for specific tasks, ultimately contributing to more effective genomic prediction models.

### Quantifying Multi-Dimensional Encoding Strategies for Non-Coding SNPs

To comprehensively evaluate encoding strategies for non-coding SNPs, we examined their performance across multiple dimensions, as summarized in Table 1 and Table 2. Our analysis revealed distinct patterns in abundance, differentiation, computational cost, and interpretability, providing insights into the trade-offs inherent in these approaches and the mechanisms behind their effectiveness.

**Table 1.**
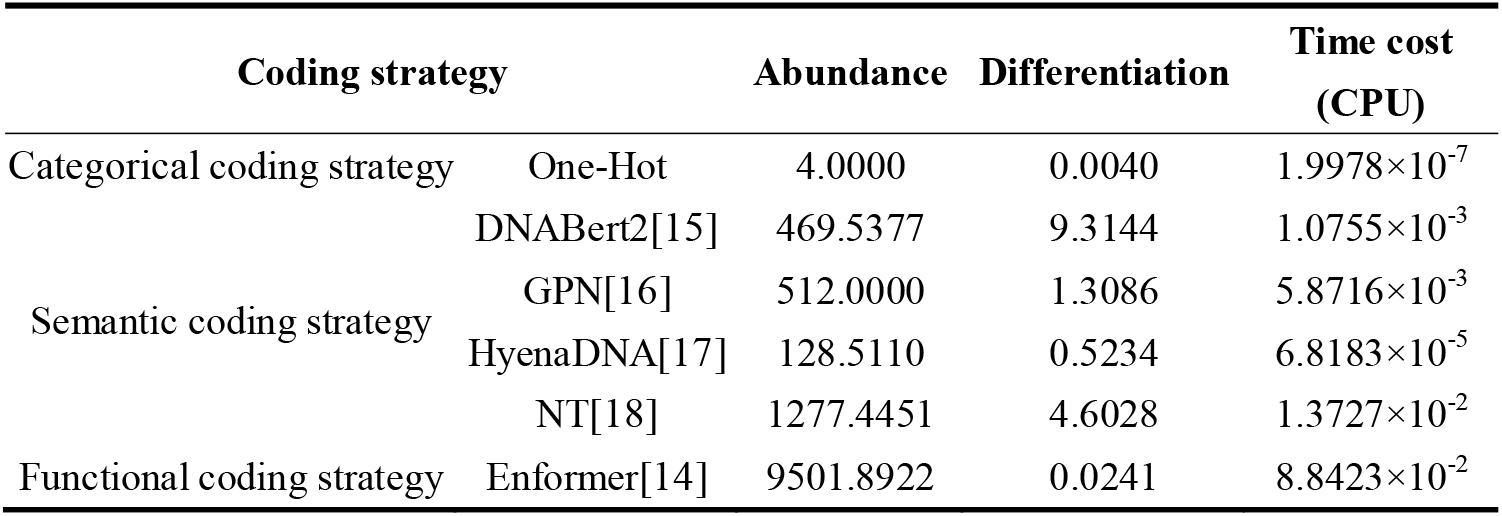
Feature profile of coding strategies.

**Table 2.**
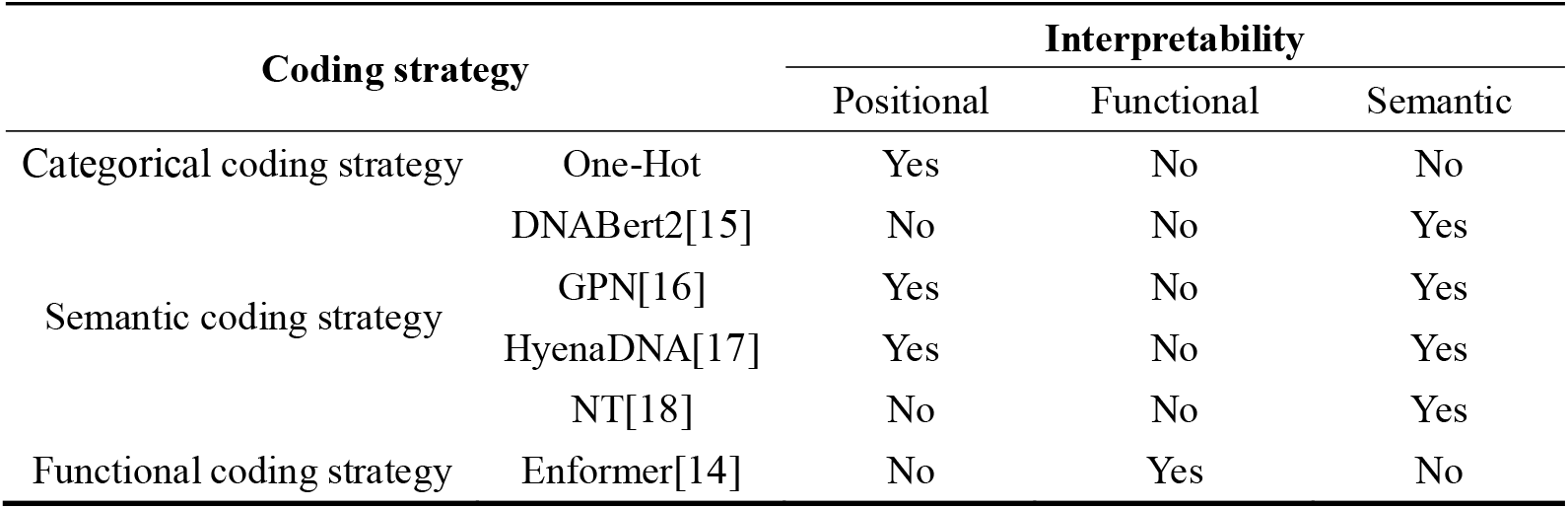
Interpretability analysis of coding strategies.

The abundance of an encoding strategy, defined as the amount of information required to describe a given mutation, varied significantly across methods. Categorical strategies such as One-Hot encoding used minimal representation dimensions (4), whereas functional strategies like Enformer required significantly more (5,313 dimensions). Semantic strategies, including DNABert2 (768 dimensions) and NT (1280 dimensions), struck a balance, offering moderately compact yet information-rich representations. Differentiation, defined as the change in embedding information per unit mutation length—quantifying the encoding’s ability to amplify the effect of mutations—exhibited notable variability across strategies. For instance, DNABert2 demonstrated the highest differentiation score (9.31), while Enformer exhibited a low score (0.0241) despite having the highest abundance. This highlights the diversity of encoding mechanisms, where higher abundance does not necessarily imply a stronger amplification of mutation effects as measured by differentiation.

Computational efficiency was another critical factor, particularly for large-scale genomic studies. One-Hot encoding was the most computationally efficient, with a negligible time cost of 1.9978×10^−7^ CPU seconds per SNP. In contrast, Enformer was the most computationally demanding at reflecting its reliance on extensive functional annotations and high-dimensional representations. Among semantic-wise strategies, DNABert2 achieved a favorable balance with a time cost of 1.0755×10^−3^, while HyenaDNA offered slightly faster embeddings at 6.8183×10^−5^.

Interpretability, a cornerstone of biological utility, was examined across three dimensions: positional, functional, and semantic. Positional interpretability refers to an encoding strategy’s ability to preserve explicit sequence positional information, enabling precise alignment of variant loci with genomic regulatory elements. This characteristic was present in One-Hot encoding as well as in certain semantic embedding methods such as GPN and HyenaDNA (Table 2), where the encoded representations maintain a base-wise correspondence with the original DNA sequence. These strategies are particularly advantageous in tasks requiring spatial resolution of variant effects.

To further explore this property, we conducted a kernel density analysis of embeddings derived from these three positionally interpretable methods, comparing their distributions across enhancer and non-enhancer regions (Figure S2). The resulting distributions revealed that these embeddings effectively retained locational information and facilitated integrative analyses with functional genomic annotations. In particular, One-Hot encoding exhibited markedly non-uniform density patterns between enhancer and non-enhancer regions. This pattern echoes the nucleotide composition biases commonly observed between functional and non-functional genomic segments [29], reinforcing its capacity to capture fine-grained, sequence-level positional variation.

Functional interpretability, defined as the incorporation of explicit genomic annotations into the representation, was uniquely achieved by Enformer, which integrates multi-modal biological context, thereby enhancing downstream functional inference. In contrast, semantic interpretability, the capacity to encode abstract and high-order biological signals, was a defining feature of semantic embedding strategies such as DNABert2, NT, and HyenaDNA. These approaches captured complex dependencies within DNA sequences, providing nuanced insights into the regulatory roles of non-coding variants beyond surface-level annotations.

These findings lead to several important conclusions. First, the relationship between abundance and differentiation is non-linear, emphasizing the importance of how information is structured within embeddings rather than the sheer dimensionality of representation. Semantic strategies like DNABert2 demonstrate this principle by achieving high differentiation with moderate abundance. Second, computational cost presents practical trade-offs, particularly for large-scale studies. While One-Hot encoding remains the fastest, its limited interpretability might restrict its utility for complex tasks. Semantic-wise strategies like DNABert2 and HyenaDNA offer practical alternatives, balancing efficiency with robust differentiation and interpretability.

Finally, no single strategy excels across all dimensions, reflecting the complementary strengths of these approaches. Functional strategies like Enformer provide unparalleled biological insights at the cost of computational efficiency, while semantic strategies capture abstract sequence relationships with less emphasis on functional annotation. This diversity underscores the potential for integrating multiple strategies to enhance both predictive performance and biological interpretability.

### Evaluating Performance in Predicting Regulatory Directions of Non-Coding SNPs on Gene Expression

The expression quantitative trait loci (eQTL) sign prediction task focuses on classifying whether non-coding SNPs positively or negatively regulate gene expression. This task provides critical insights into the functional roles of non-coding variants in transcriptional regulation[14]. To systematically evaluate performance, we designed controlled variable experiments using eQTL data from the GTEx v8 dataset[30] for three tissues: esophagus mucosa, heart left ventricle, and nerve tibial. Five machine learning models and four deep learning models were tested, paired with six encoding strategies. Additionally, we assessed the impact of data preprocessing strategies, including chromosome-based splitting versus randomized splitting of training, validation, and testing datasets.

Machine learning models, optimized via grid search, demonstrated overall superior performance compared to deep learning models. This trend suggests that deep learning does not universally outperform traditional machine learning, particularly in datasets with limited sample sizes, such as the eQTL dataset used in this study. However, the difference in performance may also reflect the relative simplicity of the task, which machine learning models can address effectively with fewer parameters (Figure 2a, Supplementary Figure S1).

**Figure 2.**
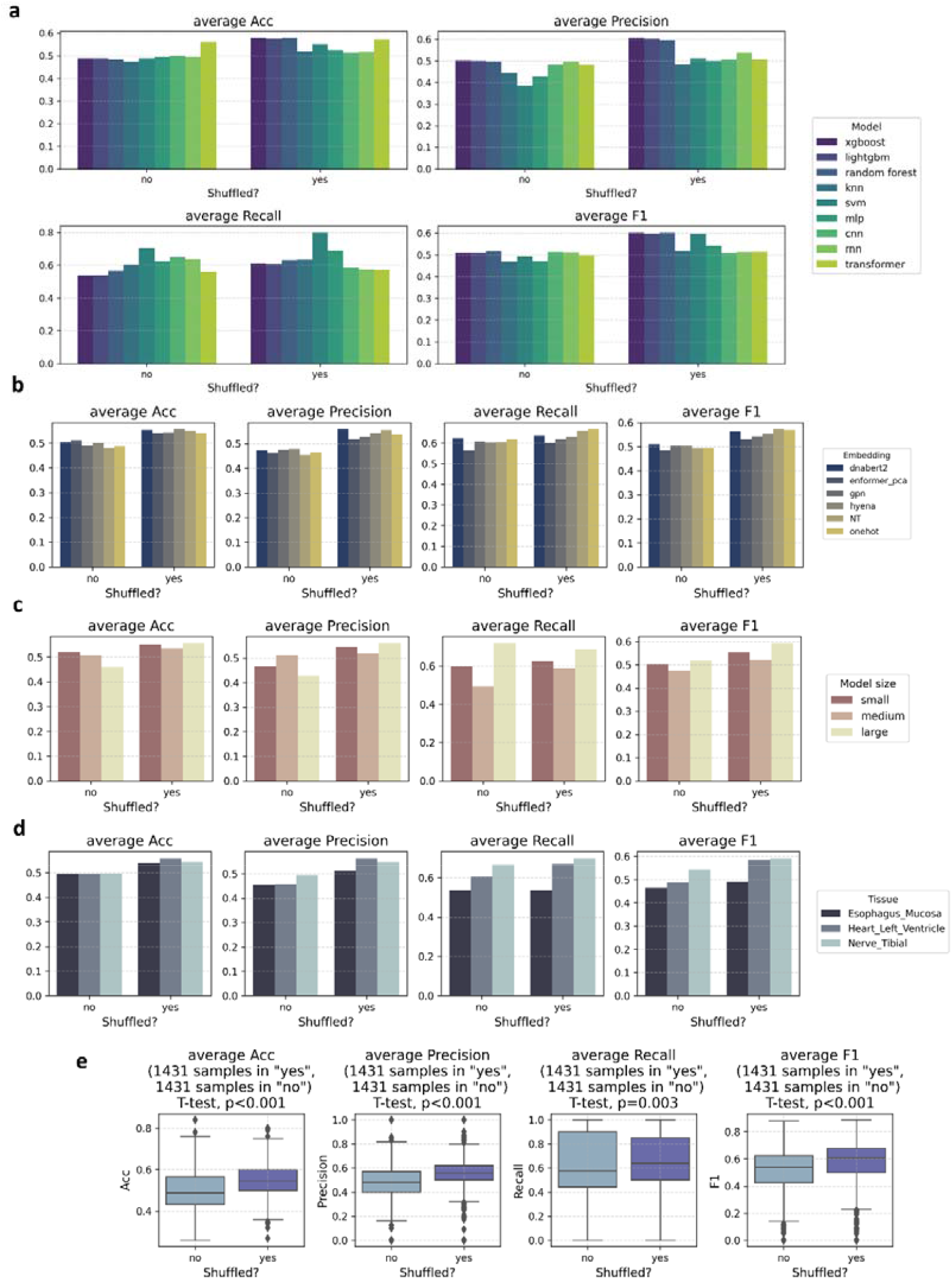
Performance analysis of eQTL sign prediction using controlled variable experiments, grouped by preprocessing strategy (“shuffled?” = “yes” indicates random splitting into training, validation, and testing sets; “shuffled?” = “no” indicates chromosome-based splitting). (a) Performance of classification tasks with the downstream machine learning/deep learning models as the primary variable. (b) Performance of classification tasks with the coding strategy as the primary variable. (c) Performance of model size (input sequence length) with the coding strategy as the primary variable. The model is divided into small (distance less than 1kbp), medium (distance between 1kbp and 10kbp), and large (distance between 10kbp and 100kbp) according to the distance from TSS to SNP. (d) Performance of classification tasks with the tissue type as the primary variable. (e) Boxplot summarizing the impact of shuffle strategies on performance. Mann–Whitney U test results indicate that differences in shuffle strategies significantly affect the performance of this downstream task.

In the evaluation of different encoding strategies, we observed minimal variation in performance across embeddings. DNABert2 and NT embeddings, despite being reduced via mean pooling, achieved performance comparable to or even slightly better than One-Hot encoding. This result highlights the potential of semantic embeddings in capturing meaningful sequence information, even after dimensionality reduction. Conversely, Enformer embeddings, which were reduced via principal component analysis (PCA)[31] to balance computational costs, did not outperform simpler encodings. This may be attributed to the inherent complexity of the Enformer model and the potential loss of information during dimensionality reduction (Figure 2b).

The regulatory distance between SNPs and the transcription start sites (TSS) emerged as another important factor. Although longer regulatory relationships are theoretically more challenging to model[32], our results did not show a consistent decline in performance with increasing sequence length. Instead, models trained on longer sequences (10-100kbp) performed comparably to those with shorter inputs. This counterintuitive observation may be explained by differences in data availability; larger datasets for longer sequences likely contributed to improved training and overall performance (Figure 2c, Supplementary Figure S1).

Performance also varied across tissues, reflecting the tissue-specific nature of gene regulation and the diversity of eQTL datasets. Differences in data distribution and regulatory complexity among tissues likely influenced these results (Figure 2d).

We further examined the impact of preprocessing strategies on model performance by comparing two commonly used data partitioning methods: chromosome-based splitting and random shuffling of eQTL datasets. Statistical analysis using the Mann–Whitney U test revealed that random shuffling was associated with higher performance across most models and encoding strategies (Figure 2e). However, this improvement does not necessarily imply that random shuffling is universally superior. The observed differences likely stem from the inherent structure of genomic data. Chromosome-based splitting preserves inter-chromosomal boundaries, thereby more closely simulating real-world generalization tasks where unseen chromosomal regions must be predicted. However, this approach may introduce distributional shifts between training and test sets, especially if regulatory elements or variant patterns are unevenly distributed across chromosomes. In contrast, random shuffling ensures that training and test data are drawn from similar sequence distributions, which may mitigate overfitting to chromosome-specific features and enhance apparent model performance.

These findings underscore the importance of aligning preprocessing choices with the intended use case. For rigorous performance evaluation and to avoid information leakage, chromosome-based splitting remains the more conservative and biologically realistic strategy. However, if the goal is to assess general model robustness or maximize predictive accuracy in a cross-genomic context, random shuffling may be appropriate. We recommend that future studies explicitly report their data splitting strategy and consider the trade-offs it entails in genomic modeling tasks.

### Evaluating Performance in Predicting Impact Magnitude of Non-Coding SNPs on Gene Expression

The task of predicting the magnitude of non-coding SNPs’ impact on gene expression is a regression problem that aims to estimate the degree of gene expression change associated with specific variants. We applied the same experimental setup and data as in the previous eQTL sign prediction task, including the GTEx eQTL datasets from esophagus mucosa, heart left ventricle, and nerve tibial tissues. Similar to the classification task, machine learning models, especially those optimized through grid search, outperformed deep learning models (except for KNN, Supplementary Figure S3). Dimensionality reduction techniques, such as PCA for Enformer and mean pooling for DNA pre-trained language models, were used to manage computational costs.

The overall performance trends were consistent with those observed in the sign prediction task (Supplementary Figure S3). The impact of encoding strategies on performance remained minimal, with semantic embeddings like DNABert2 and NT showing potential despite dimensionality reduction. Additionally, the SNP-TSS distance did not significantly affect model performance, likely due to the larger data volumes available for longer sequences. As with the previous task, random shuffling of datasets significantly enhanced performance, underscoring its importance for robust training and evaluation. These results confirm the robustness of our findings from the classification task, emphasizing the effect of shuffle strategies and highlighting the potential of semantic encoding strategies.

### Evaluating Performance in Predicting Impact Magnitude of Non-Coding SNPs on DNA Methylation Levels

We then focus on evaluating the performance of different coding strategies in predicting the impact magnitude of non-coding SNPs on DNA methylation levels, specifically through meQTL slope prediction. For this task, we utilized the meQTL EPIC dataset[33], which includes CD4+ T cells and monocytes. As with the eQTL slope prediction, the overall performance trends follow similar patterns across different models and strategies.

Machine learning models, particularly those optimized via grid search, continued to perform robustly across most settings except for KNN (Supplementary Figure S4). Notably, the performance gap between machine learning and deep learning models was smaller in this task, potentially reflecting the stabilizing effect of a larger dataset. This suggests that while machine learning models offer consistent performance across varying conditions, deep learning architectures may hold greater potential under settings with higher data volume and increased regulatory complexity.

In terms of encoding strategies, Enformer embeddings—despite dimensionality reduction via PCA—achieved the highest Pearson correlation in some scenarios, suggesting that biologically annotated representations may retain advantageous functional signals for methylation prediction (Figure 2f). At the same time, semantic embeddings such as DNABert2 and NT remained competitive, even after mean pooling, highlighting their robustness and adaptability across downstream tasks. The variation in model performance across cell types further reflects the diversity of SNP–CpG interactions and the cell-type-specific regulatory mechanisms embedded in the meQTL landscape.

### Univariate Analysis: Investigating the Relationship Between Model Complexity, Tissue Type, and Performance

In QTL prediction tasks, it is widely believed that longer genomic regions, such as those with greater TSS-to-variant distances, may result in more complex regulatory relationships and, consequently, reduced model performance. However, our experiments led to an unexpected observation in the eQTL sign prediction task. Despite the intuitive expectation that larger models, which consider more extended genomic regions, would perform worse, we found that the large model outperformed the middle model (Figures 3a-b). This suggests that, contrary to prior assumptions, other factors may play a critical role in shaping model performance. Upon further inspection, we observed that the large model indeed utilized a greater sample size than the middle model (Supplementary Figure 1c), which points to the possibility that sample size, an initially unconsidered variable, could be influencing the model’s ability to effectively fit the data. Inadequate data volume may hinder the model’s capacity to capture the full complexity of the genomic signals, limiting its predictive accuracy.

**Figure 3.**
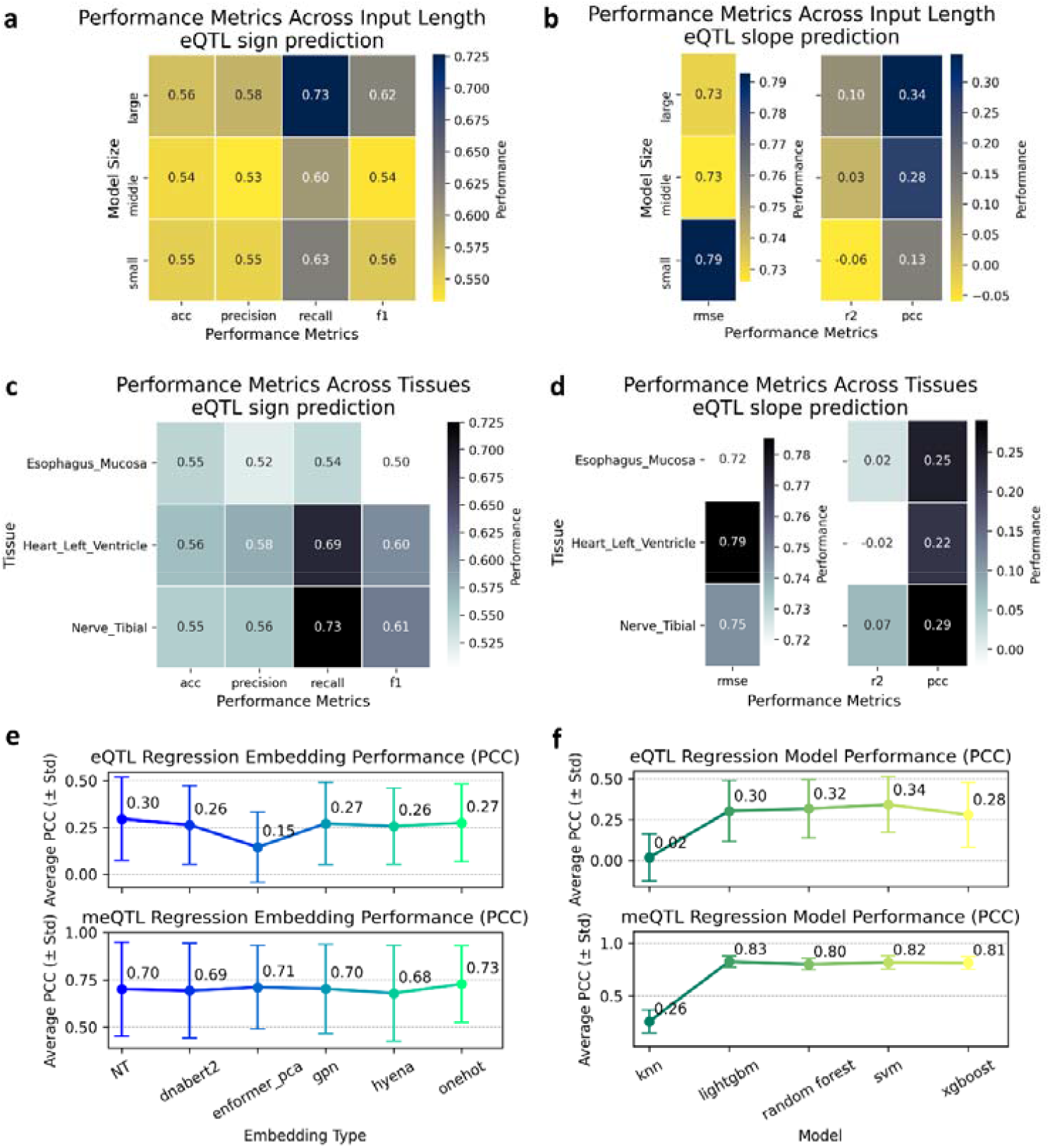
Univariate Analysis of QTL Prediction Tasks. (a)-(d) Investigation of the model performance with model size and tissue type as observed explicit variables and corresponding sample size as the implicit variable. (e)-(f) Examination of the relationship between embeddings, downstream task modeling strategies (machine learning), and model performance on slope prediction tasks.

To further explore this, we examined whether the performance differences could be attributed to sample size or the inherent differences between tissue types. Specifically, we compared the data volumes used for training across various tissues (Supplementary Figure 1a) and found that the heart left ventricle tissue, with the smallest dataset, did not necessarily exhibit the worst model performance (Figure 3c-d). This suggests that while sample size may have an impact, tissue type itself could also be influencing model performance. The variation in regulatory mechanisms across tissues likely leads to differences in fitting difficulty and generalization, highlighting the complex interplay between sample size, tissue type, and prediction performance.

### Cross-QTL Task Analysis: Exploring the Effectiveness of Encoding Strategies and Models

To further understand the impact of encoding strategies and model choices on QTL prediction tasks, we conducted cross-QTL task analysis by examining the effectiveness of different embeddings in slope prediction (Figure 3e-f). In this experiment, we compared the performance of various encoding strategies, across two regression tasks. Interestingly, the performance differences between encoding strategies in the meQTL slope prediction task were relatively minor. Given the large sample size of the meQTL dataset, it is likely that all embedding strategies were sufficiently powerful to capture the underlying regulatory patterns, resulting in a comparable model performance (Figure S1).

In contrast, when applied to the eQTL tasks, performance discrepancies were more evident, particularly when Enformer was reduced via PCA to fit the task’s requirements. This suggests that dimensionality reduction, while necessary to handle the high computational cost of Enformer, may hinder its effectiveness in tasks with smaller sample sizes, such as eQTL. In terms of modeling approaches, tree-based algorithms (XGBoost, LightGBM, and Random Forest) performed consistently well across the tasks. SVM showed similar performance to the tree models, but the KNN algorithm underperformed. KNN’s poor performance in QTL prediction tasks can be attributed to its reliance on local similarity and inability to capture the complex, non-linear relationships and feature interactions present in high-dimensional genomic data.

### Guided Analysis for Practical Use: Case Study on meQTL Prediction

In the case study, we aimed to offer a guided analysis for practical use in meQTL prediction, providing a comprehensive framework for how one might approach QTL-related tasks. This study is not only a showcase of the potential of our framework but also serves as a reference for others looking to implement QTL prediction with genomic data. We began by visualizing the data distribution for monocyte meQTL, which provided an intuitive overview of the dataset, helping contextualize the performance of subsequent analysis (Figure 4a). For the performance evaluation, we employed a regression task using PCA-reduced Enformer embeddings as input and a Transformer framework for representation learning. This allowed us to assess the model’s predictive power on the test set (Figure 4b).

**Figure 4.**
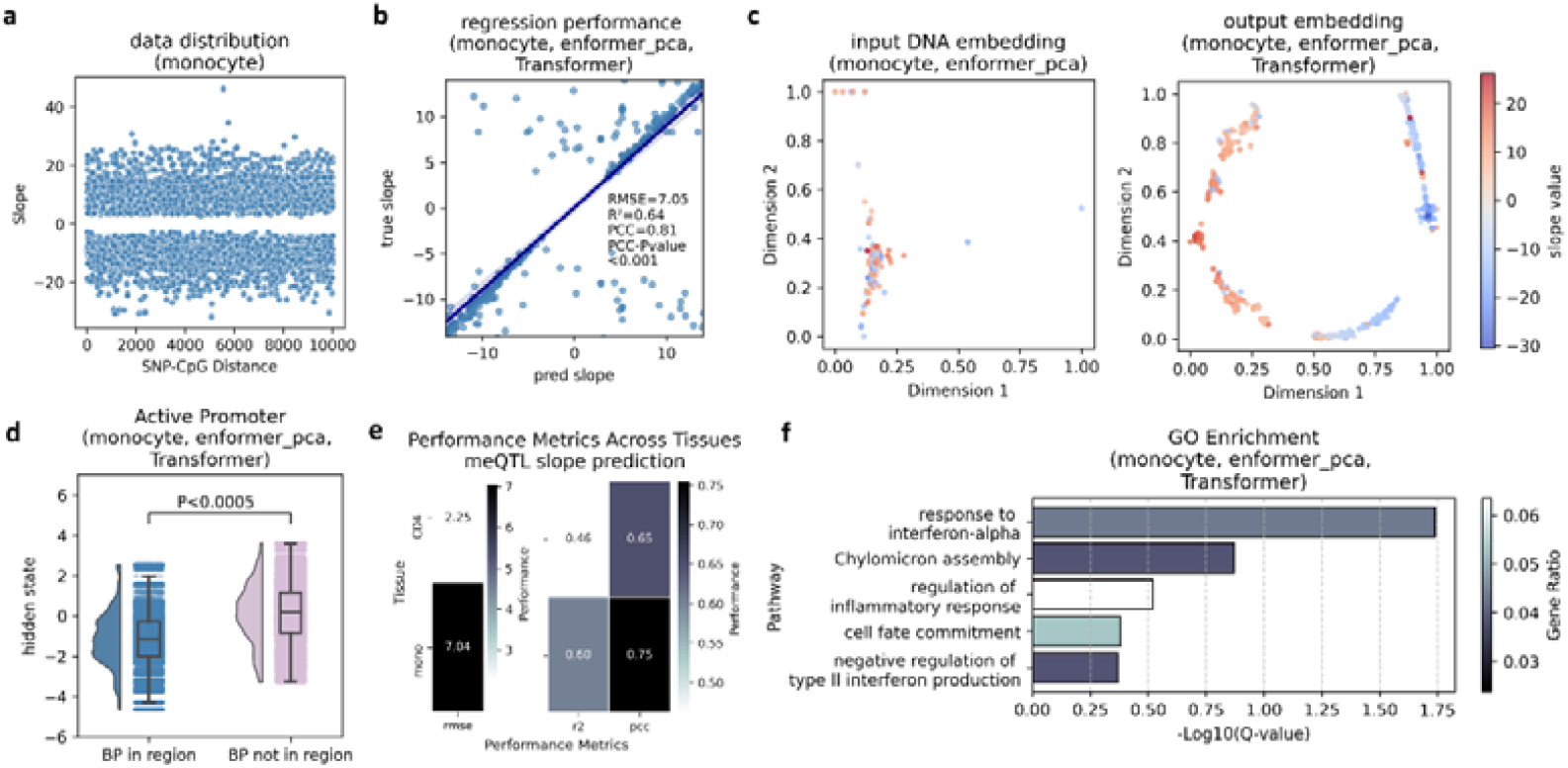
Case study on meQTL prediction as a guided analysis for practical use. (a) Visualization of the data distribution for monocyte meQTL to provide an intuitive overview of the dataset. (b) Performance evaluation of a regression task on the test set using PCA-reduced Enformer embeddings as input and a Transformer framework for representation learning. (c) Visualization of embeddings before and after model input using t-SNE dimensionality reduction, showcasing the model’s representation learning capability. (d) Analysis of the model’s interpretability by annotating hidden states corresponding to bases located in active promoter regions. (e) Evaluation of model performance across different cell types, with data volume as the implicit variable. (f) GO enrichment analysis of genes associated with SNPs predicted to have high impacts (predicted slope absolute value > 0.2), uncovering potential links to disease mechanisms.

Further investigation into the model’s behavior was conducted by visualizing the embeddings before and after model input with t-SNE dimensionality reduction, revealing how well the model learned meaningful representations of the data (Figure 4c). We also performed an interpretability analysis by annotating hidden states corresponding to bases located in active promoter regions, highlighting the model’s ability to link genomic features with regulatory functions (Figure 4d). Additionally, we evaluated model performance across different cell types, observing how varying data volumes influenced the results (Figure 4e). Lastly, a Gene Ontology (GO) enrichment analysis[34] of genes associated with SNPs predicted to have high impacts (slope absolute value > 0.2) unveiled potential links to disease mechanisms, offering valuable insights for future investigations (Figure 4f). This case study underscores how such analyses can be integrated to optimize QTL prediction tasks.

## Discussion

In this study, we explored and quantified the performance of various encoding strategies and models in predicting the regulatory effects of non-coding SNPs. By evaluating multiple encoding strategies, including categorical, semantic, and functional embeddings, across several downstream tasks, we identified the effectiveness and potential trade-offs associated with each strategy in different genomic contexts. Focusing on tasks such as eQTL and meQTL prediction, we employed a controlled variable approach to ensure the robustness and reliability of our findings. Our comparison of machine learning and deep learning models further highlighted the nuanced interplay between model choice and data characteristics, such as sample size and machine/deep learning models. Moreover, the analysis of preprocessing strategies emphasized the critical role of dataset splitting in ensuring accurate model evaluation. This work provides actionable insights for advancing the analysis of non-coding SNPs, ultimately contributing to a deeper understanding of their biological roles.

Our results highlight the multifaceted nature of predictive performance, shaped by interactions among encoding strategy, model architecture, sample size, and data preprocessing. Machine learning models, particularly when optimized via grid search, consistently delivered stable performance, especially in moderately sized datasets. Meanwhile, semantic embeddings demonstrated strong potential in capturing regulatory patterns, even after dimensionality reduction, indicating their robustness across diverse settings.

We also examined the impact of data splitting strategies on model evaluation outcomes. While random shuffling and chromosome-based partitioning each yielded different trends in performance, neither approach emerged as universally superior. Instead, the observed differences underscore the importance of selecting preprocessing strategies based on the specific goals of a given study. For instance, chromosome-based splitting may better reflect generalization across genomic regions, whereas random shuffling can be useful for assessing performance consistency under distributionally uniform conditions.

Importantly, the performance of these encoding strategies was found to be relatively consistent across different QTL prediction tasks, indicating that once embedded representations are fully learned, they effectively capture the regulatory information embedded within the genomic sequence. The study also revealed that sample size might also be crucial in model performance, particularly for smaller datasets, where model fitting may not reach optimal levels. Additionally, the cross-QTL task analysis underscored the importance of choosing appropriate models, with tree-based machine learning models like XGBoost and LightGBM outperforming KNN, which may struggle to capture complex relationships in high-dimensional genomic data.

The case study on meQTL prediction further demonstrated the practical utility of the methods developed in this work, providing actionable insights for researchers in the field. Our guided analysis also exemplified how the interpretability of models can be enhanced through visualizations such as t-SNE and by annotating hidden states, offering a more comprehensive understanding of SNP impacts on methylation. This comprehensive approach not only contributes to advancing meQTL prediction but also sets the stage for future work aimed at refining predictive models, integrating additional genomic information, and scaling these methods to broader datasets.

While this benchmark provides valuable insights into the comparative performance of various encoding strategies and models, several limitations remain. First, the study primarily focuses on a relatively small set of cell types and tissues, and extending the analysis to a broader range of cellular contexts would allow for more generalized conclusions. Additionally, while we have assessed a variety of encoding methods and models, there may exist other, more optimal combinations or novel encoding approaches that were not included in this study, warranting further exploration. Future work could also focus on improving the performance of models on small sample sizes and enhancing their generalization capabilities. Ultimately, this benchmark establishes a foundational framework for future studies on non-coding SNPs, offering actionable insights for the design of predictive pipelines and the selection of appropriate models for specific genomic tasks.

## Materials and Methods

### Datasets

For eQTL data, we downloaded eQTL for three tissues: esophagus mucosa, heart left ventricle, and nerve tibial from the GTEx v8 dataset[30]. These datasets were used to evaluate the regulatory effects of non-coding SNPs on gene expression across diverse genomic contexts. For meQTL data, we utilized the meQTL EPIC dataset, which included meQTL data for two cell types: CD4+ T cells and monocytes. The distributions of these datasets are visualized in Supplementary Figure S1.

In this study, eQTL datasets were annotated and DNA sequences were retrieved based on the GRCh38/hg38 reference genome, while meQTL datasets were processed using the GRCh37/hg19 reference genome. Genomic sequences and their functional annotations were obtained from the UCSC Genome Browser, ensuring consistency in annotation and feature extraction.

For datasets without random shuffling, we used a chromosome-based splitting strategy during model training. Specifically, the training set consisted of data from chromosomes 1-9 and 14-22, the validation set was derived from chromosomes 12-13, and the test set was sourced from chromosomes 10-11.

### Coding strategies

In this study, all encoding strategies require DNA sequence sampling, with the sampling standard differing between tasks. For eQTL prediction tasks, the sequence is sampled with the TSS at the center, ensuring the mutation is within the sampling range. For meQTL prediction tasks, the sequence is sampled with the CpG site at the center, ensuring the mutation is included in the sampling range. Specifically, for eQTL prediction, sequence lengths of 2,001bp, 20,001bp, and 200,001bp are used, while for meQTL tasks, a sampling length of 20,001bp is applied.

- **OneHot**. The DNA sequence is represented by a one-hot encoded vector with four possible values corresponding to the four bases (A, T, C, G). If a base is ‘N’ (unknown), it is encoded as four zeros.
- **DNABert2[15]**. DNABert2 is a pre-trained DNA language model, which was implemented using the official API (https://github.com/MAGICS-LAB/DNABERT_2) in this study. For DNABert2, we designed a position-wise average cutting and pooling strategy, aiming to maximize mutation representation. The processing steps are as follows: given a DNA sequence centered on either the TSS (for eQTL) or CpG site (for meQTL), the two outer segments are cut into lengths of 250bp, while the center segment (501bp) is centered around the TSS or CpG site. The remaining parts of the DNA sequence between the two outer segments and the central segment are evenly divided into 500bp fragments. For each fragment, an embedding is generated, and embeddings are concatenated along the embedding dimension. Finally, the concatenated embeddings are averaged over the DNA sequence dimension to produce a one-dimensional vector of the same dimension as the language model embedding.
- **GPN[16]**. GPN is a pre-trained DNA language model, implemented using the official API (https://github.com/songlab-cal/gpn) in this study. The same position-wise average cutting and pooling strategy applied to DNABert2 is also used here.
- **HyenaDNA[17]**. HyenaDNA is another pre-trained DNA language model, implemented via the official API (https://github.com/HazyResearch/hyena-dna) in this study. Like DNABert2, it applies the position-wise average cutting and pooling strategy.
- **NT[18]**. NT is a pre-trained DNA language model, implemented using the official API (https://github.com/instadeepai/nucleotide-transformer) in this study. It also utilizes the position-wise average cutting and pooling strategy, similar to DNABert2.
- **Enformer[14]**. Enformer is a model designed for gene expression prediction from DNA sequences. In this study, we extracted DNA sequences of length 196,608 bp centered around the TSS or CpG sites to match Enformer’s input requirements. For both pre-mutated and post-mutated sequences, we obtained output features with a dimension of 5,313. When calculating quantitative metrics, we directly utilized these output features. However, to balance computational cost during downstream task validation, we performed PCA dimensionality reduction on these features, retaining the top 10 principal components. The use of PCA for Enformer representations was motivated by the model’s architectural design: each dimension in its output corresponds to a biologically interpretable feature, such as specific regulatory annotations. Applying PCA in this context enables the extraction of the most informative components while preserving biologically relevant variation, potentially facilitating the discovery of core regulatory signals.

### Feature profile of coding strategies

To comprehensively evaluate the strengths and limitations of different encoding strategies, we defined a set of feature profiles to quantify their capabilities in representing non-coding SNPs.

- **Abundance**. Abundance measures the amount of information required to describe a mutation. Specifically, it refers to the dimensionality of the embedding per base, indicating how detailed the representation is for each SNP. The final value is normalized by the sequence length *L*, measured as the length of the DNA sequence centered on the mutation:

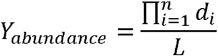

where *d*_i_ is the dimensionality of the *i*-th embedding axis.
- **Differentiation**. Differentiation quantifies the change in embedding information per unit length of DNA before and after mutation, effectively capturing the ability of an encoding strategy to amplify mutation effects. This difference is computed for each dimension of the embedding vector, *d*_i_, and averaged over the entire sequence length *L* centered around the mutation. The formula can be expressed as:

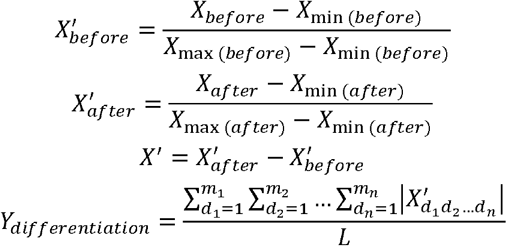

where *X*′represents the normalized difference in the *i*-th dimension of the embedding vector before and after mutation.
- **Time cost**. Time cost represents the computational expense required to generate embeddings for a unit length of DNA, both before and after mutation. This metric reflects the efficiency of the encoding strategy, which is particularly critical for large-scale genomic datasets:

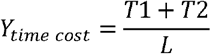

where *T*1 and *T*2 represent the computational time required to generate the embedding vectors for the unmutated and mutated DNA sequences, respectively.
- **Interpretability**. Interpretability assesses whether the encoding preserves key biological information, such as the relative positions of bases or functional annotations of the DNA sequence. This dimension is crucial for understanding the biological implications of predictions made by models using these encodings.

(1) **Positional Interpretability**: the clarity with which the embedding can be directly mapped to individual nucleotides in the DNA sequence. This means the position of each nucleotide in the original sequence can be unambiguously represented in the embedding.
(2) **Functional Interpretability**: whether the embedding captures biological features that can be derived from experimental techniques such as sequencing. This includes elements like transcription factor binding sites, enhancers, or other functional genomic elements that are detectable through high-throughput sequencing methods.
(3) **Semantic Interpretability**: the contextual meaning of the sequence as learned through pre-training models. It reflects the ability of the embedding to encapsulate not just the individual bases, but also the broader biological context in which these sequences operate, as learned from large-scale sequence data.

In this work, we do not investigate the biological properties that might be encoded in embeddings derived solely from pre-trained DNA sequence models, as these properties are currently not directly modelable in this context.

### Machine learning and deep learning models

For machine learning models, we selected eXtreme Gradient Boosting (XGBoost)[21], Light Gradient Boosting Machine (LightGBM)[22], Random Forest (RF)[23], k-Nearest Neighbors (KNN)[24], and Support Vector Machine (SVM)[25]. To optimize these models, we utilized the GridSearchCV function from the Scikit-learn Python library, which systematically searches for the best combination of hyperparameters to achieve locally optimal solutions.

For deep learning models, we explored Multi-Layer Perceptrons (MLP)[26], Convolutional Neural Networks (CNN)[26], Recurrent Neural Networks (RNN)[27], and Transformer-based architectures[28]. These frameworks were chosen to evaluate their capacity for representation learning and their ability to model complex relationships within the genomic data.

To ensure the robustness of our findings, all experiments were conducted with three different random seeds, reducing variability and enhancing the reliability of our conclusions. This systematic approach allowed for a comprehensive comparison between machine learning and deep learning models in the context of QTL prediction tasks.

### Performance evaluation

For the classification task, we employed Accuracy (ACC), Precision, Recall, F1-score, and Area Under the Curve (AUC) to assess model performance. For the regression task, we utilized Root Mean Square Error (RMSE) and Pearson Correlation Coefficient (PCC) as key metrics.

## Data availability

The tissue-specific eQTL data were collected from the GTEx v8 portal[30] (https://www.gtexportal.org/home/datasets). The meQTL EPIC dataset[33] was downloaded from the meQTL EPIC Database website (https://epicmeqtl.kcl.ac.uk/). The GRCh37/hg19 genome, GRCh38/hg38 genome, and functional annotation were obtained from the UCSC Genome Browser (https://genome.ucsc.edu/). The CpG annotation of Infinium MethylationEPIC v1.0 B5 manifest file was downloaded from https://support.illumina.com/downloads/infinium-methylationepic-v1-0-product-files.html.

## Code availability

The source codes of the pre-processing, modeling, and validation processes are freely available on GitHub (https://github.com/Liuzhe30/DNAMutBenchMark).

## Acknowledgments

This work was supported by grants from STI 2030—Major Projects (no. 2022ZD0209100) and the Medical-Engineering Cross Foundation of Shanghai Jiao Tong University (No. YG2022ZD026, 23×010302269).

## Key points

- We systematically benchmarked six DNA sequence-based encoding strategies, including categorical, functional, and semantic embeddings, across key QTL prediction tasks.
- We highlighted the importance of shuffle strategy and sample size, showing how to better generalize and capture underlying regulatory patterns in QTL data.
- We demonstrated the robustness of semantic embeddings, such as DNABert2 and NT, even after dimensionality reduction, across diverse genomic prediction tasks.
- We proposed a practical framework integrating interpretability analyses and pathway enrichment, providing actionable guidance for designing predictive pipelines for non-coding SNPs.

