## Supplementary Materials for "Benchmarking Encoding Strategies for Non-Coding Mutations in Sequence-Based Prediction Tasks"

**Supplementary Information**

### Figures

#### Figure S1. The data distribution of the benchmark datasets in pre-processing progress.

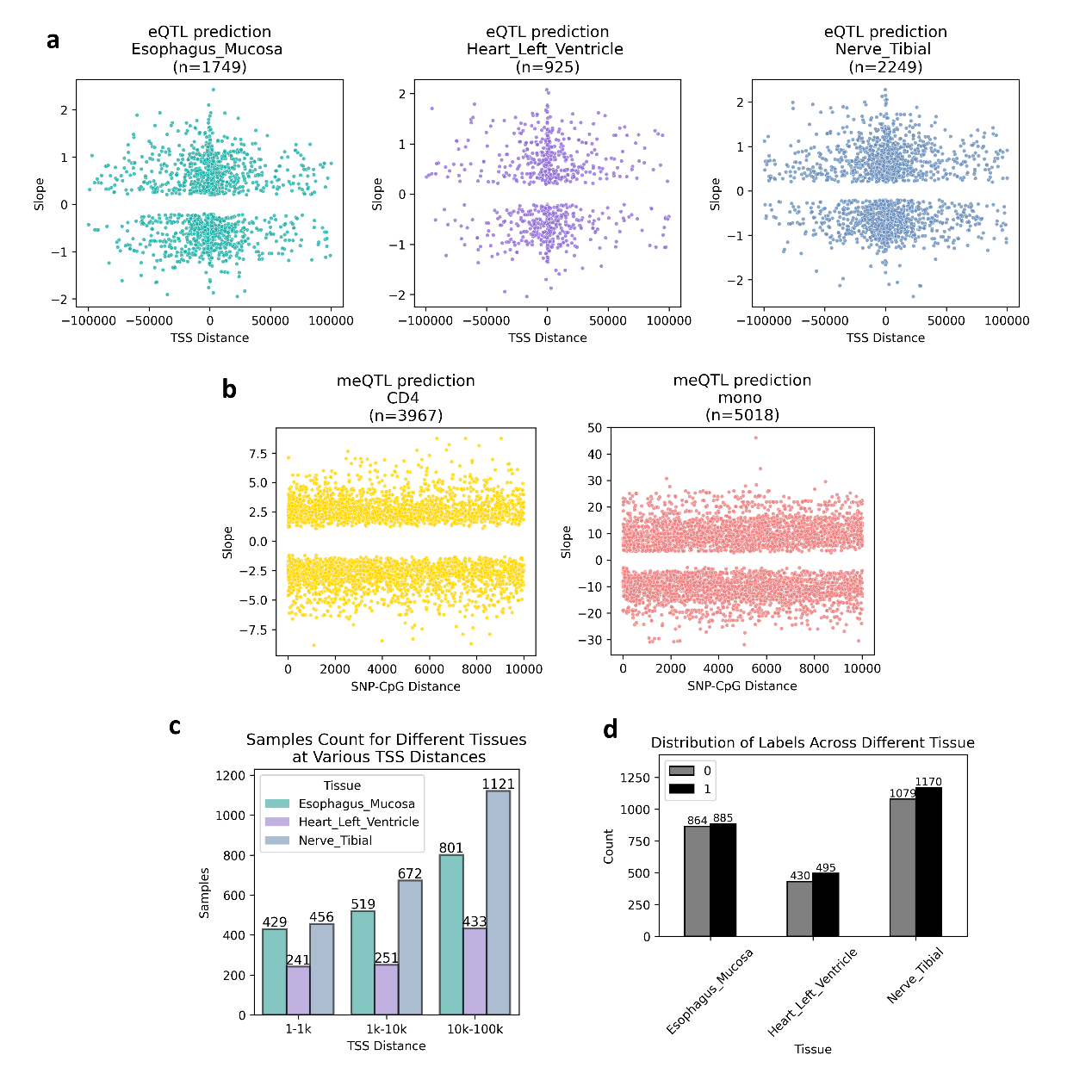

#### Figure S2. Kernel density plots of three positionally interpretable coding strategies in enhancer and non-enhancer regions. (a) Kernel density plot of GPN embeddings for enhancer and non-enhancer regions, illustrated using three example SNPs. (b) Kernel density plot of One-Hot embeddings for enhancer and non-enhancer regions, illustrated using three example SNPs. (c) Kernel density plot of Hyena embeddings for enhancer and non-enhancer regions, illustrated using three example SNPs.

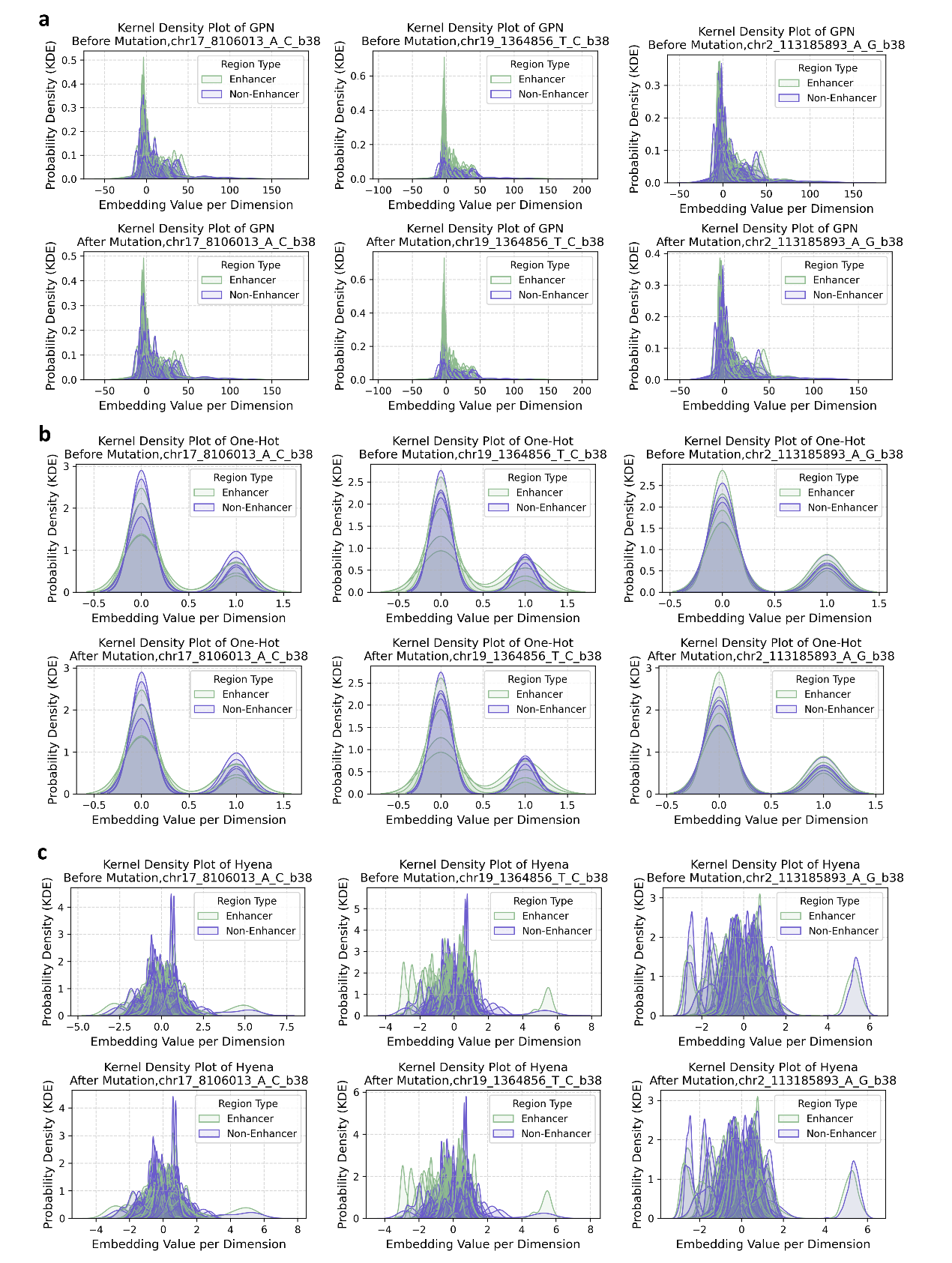

#### Figure S3. Performance analysis of eQTL slope prediction using controlled variable experiments, grouped by preprocessing strategy ("shuffled?" = "yes" indicates random splitting into training, validation, and testing sets; "shuffled?" = "no" indicates chromosome-based splitting).

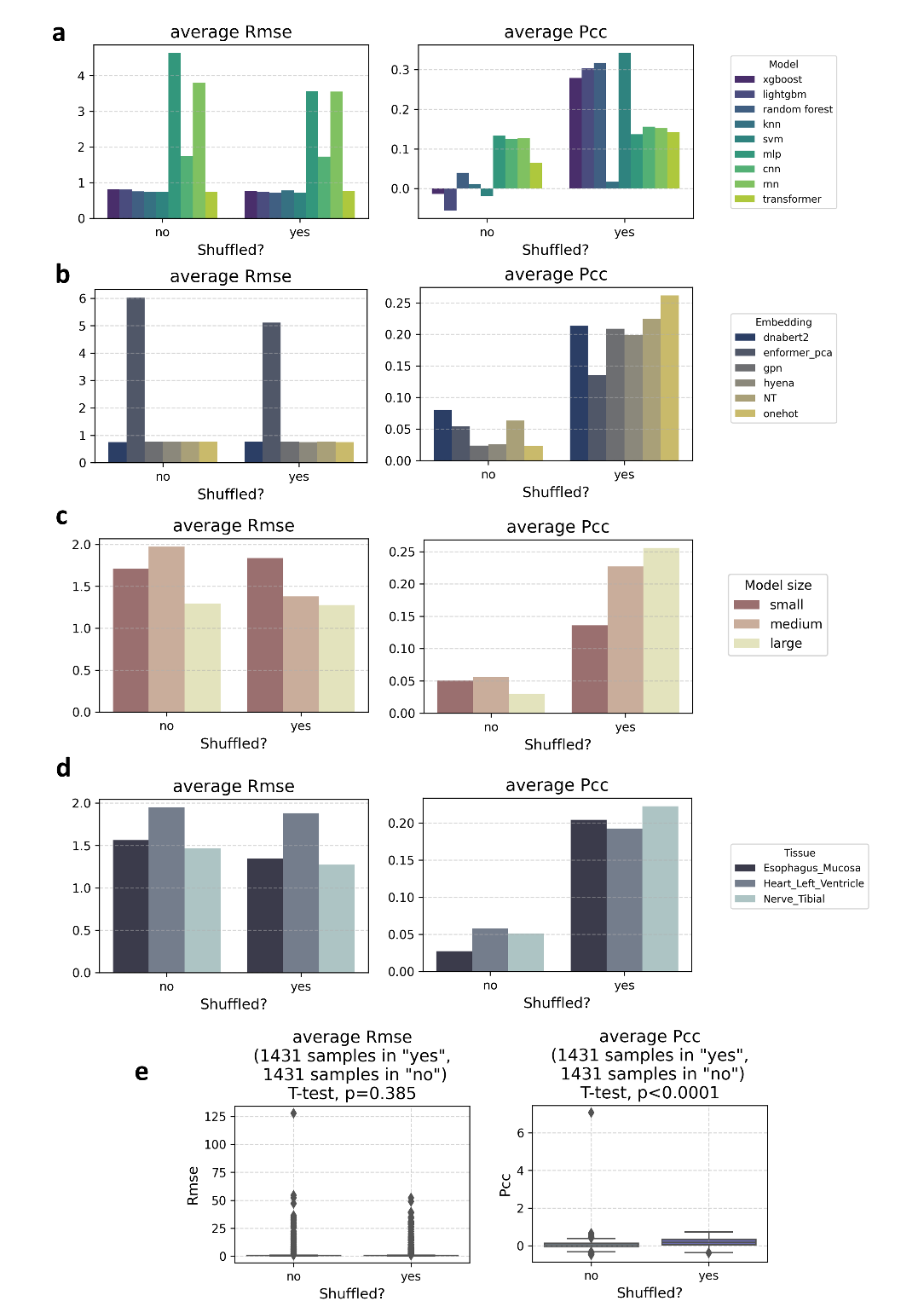

#### Figure S4. Performance analysis of meQTL slope prediction using controlled variable experiments, grouped by preprocessing strategy ("shuffled?" = "yes" indicates random splitting into training, validation, and testing sets; "shuffled?" = "no" indicates chromosome-based splitting).

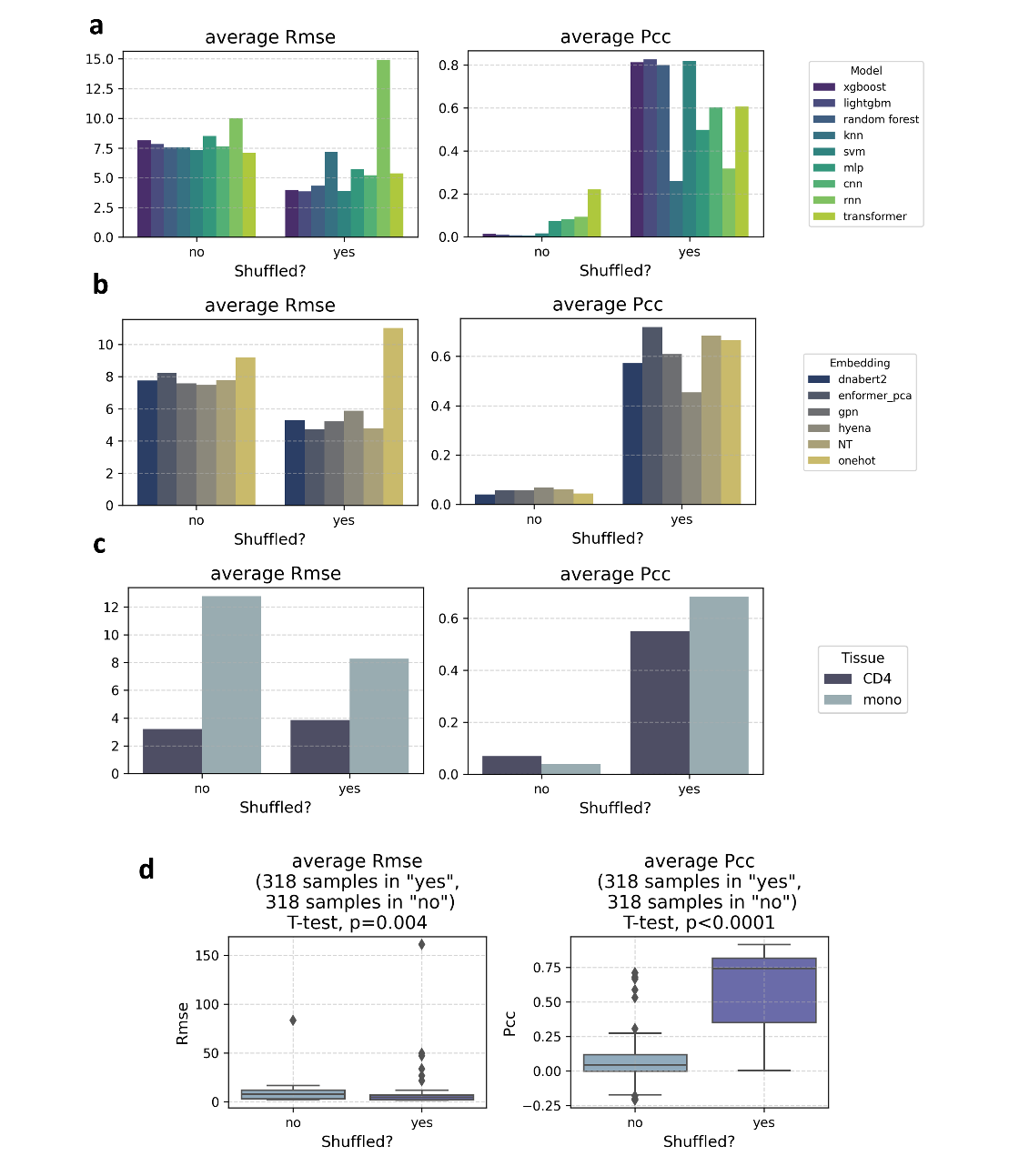

#### Figure S5. Investigation of the relationship between sample size and model performance, with model size and cell type as observed explicit variables and corresponding sample size as the implicit variable.

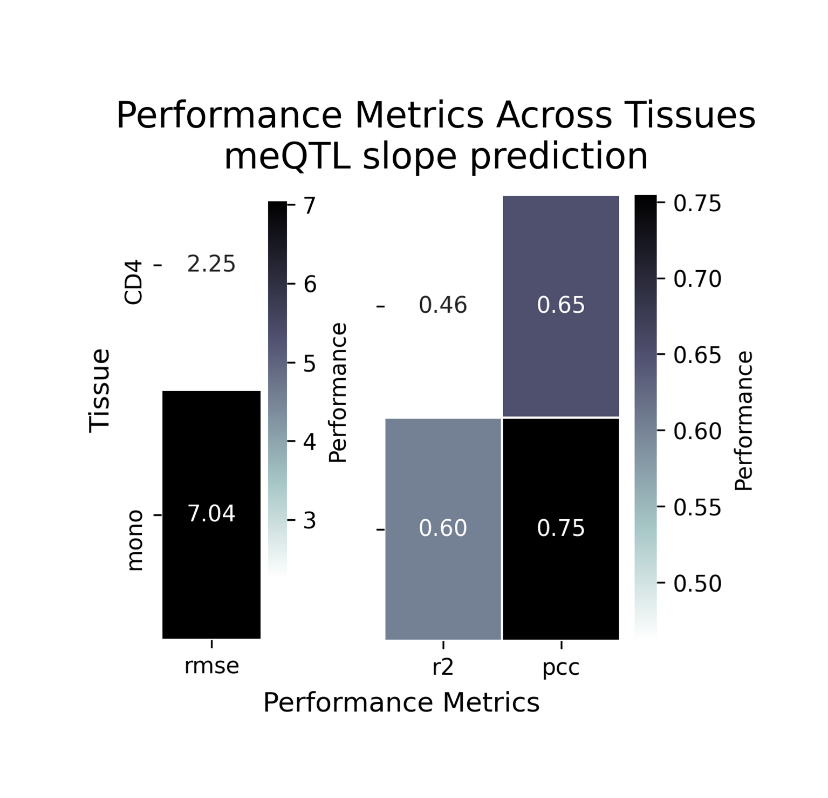
